# Loss of *asgr1a* leads to the secretion of excess dietary cholesterol in zebrafish

**DOI:** 10.1101/2024.06.26.600848

**Authors:** Tabea O.C. Moll, Joshua T. Derrick, Darby W. Sweeney, Jeffrey Shin, Steven A. Farber

**Author notes:** Corresponding author Email: Johns Hopkins University, Department of Biology 3520 San Martin Dr., Baltimore, MD 21218. The authors have declared that no conflict of interest exists.

## Abstract

One of the major pathways to clear glycoproteins from circulation is via the liver-specific asialoglycoprotein receptor (ASGPR). Loss of asialoglycoprotein receptor 1 (*ASGR1*), the major subunit of ASGPR, was recently found to correlate with lower levels of plasma apolipoprotein B-containing lipoproteins (B-lps) and a profoundly reduced risk of cardiovascular disease in humans. We set out to identify the zebrafish ortholog of *ASGR1* (*asgr1a*) and generated two independent mutations in *asgr1a* using CRISPR/Cas9. Neither *asgr1a* mutation displayed changes in larval, juvenile, and adult B-lp numbers or sizes. However, when challenged with a Western diet, *asgr1a* mutant zebrafish exhibit less hepatic steatosis and lower hepatic triglyceride levels compared to control animals. Instead, the excess dietary cholesterol was excreted. While these results do not explain the cardioprotective nature of ASGR1 in humans, they indicate the importance of ASGR1 in modulating whole animal cholesterol flux.

## Introduction

The asialoglycoprotein receptor (ASGPR) is a well-known liver-specific receptor, discovered in the 1980’s by Gilbert Ashwell and Anatol Morell (1), that is also referred to as the Ashwell receptor. The ASGPR was initially described to explicitly bind circulating proteins without terminal-sial groups (1, 2). After the removal of the sial-groups, glycoproteins that present with terminal galactose or N-acetylgalactosamine (GalNAc) groups are bound by the ASGRP (3). Once a ligand is bound, the ASGPR undergoes clathrin-mediated endocytosis. After dissociation of the ASGPR with its ligand, ASGPR is returned to the cell surface, and the ligand undergoes degradation through the lysosomal pathway (4).

ASGPR is grouped into the larger class of C-type lectin proteins (CLECs), with its two subunits, ASGR1 and ASGR2, being the only two members of the CLEC4H family. Its closest relative is the CLEC10A receptor (for a recent review, see (5)). Both ASGR1 and ASGR2 contain a cytoplasmic domain with an endocytosis motif, a transmembrane domain, followed by the extracellular carbohydrate recognition domain, the C-type lectin domain (6). Though similar in sequence, ASGR1 is required for ASGPR function, while ASGR2 is dispensable (7, 8). Changes in ligand specificity can be achieved by various forms of assembly of the ASGPR (9), which can be built by multiple possible dimers, trimers, and tetramers of the two subunits ASGR1 and ASGR2 (6, 9, 10). This makes for a wide range of ligands, including apoptotic liver cells (11), circulating tumor cells (12, 13), viruses such as SARS-CoV-2 (14–17), and lipoproteins (18, 19).

Lipoproteins are crucial for transporting hydrophobic lipids and fat-soluble nutrients through the circulation to peripheral tissues (20). Apolipoprotein B (APOB)-containing lipoproteins (B-lps) are produced by the intestine (chylomicrons) and the liver (Very-Low-Density Lipoproteins, VLDL) to transport dietary and endogenous lipids (21, 22). APOB is a required, non-interchangeable scaffold protein for B-lps (23), that remains associated with the B-lp from its production in the ER (24) to lysosomal degradation after the uptake of the B-lp from circulation (25). In circulation, lipases release the triglycerides and other contents of B-lps, effectively shrinking B-lps in size, leading to chylomicron remnants and Low-Density-Lipoproteins (LDL), respectively. Several overlapping mechanisms are responsible for the liver uptake of chylomicron remnants and LDL (19, 26, 27). The predominant uptake method is through the LDL receptor (LDLR) (27). However, since both types of particles remain in circulation for a prolonged time, they can become de-sialated and thus become ligands for the ASGPR (19).

Elevated levels of LDL are associated with an increased risk for cardiovascular disease (28). Various drugs, such as statins (29) and proprotein convertase subtilisin/kexin type 9 (PCSK9) inhibitors (30), aim to increase surface levels of LDLR to lower plasma LDL. In 2016, a genome-wide association study found that heterozygous loss of *ASGR1* in humans correlated with reduced plasma LDL levels and decreased CVD risk (31). Cell culture experiments mimicking the human 12 base pair (bp) deletion showed that the mutation reduced protein levels of *ASGR1*. Since the ASGPR is also known to bind and endocytose plasma LDL, these results were unexpected (19).

Recent work in HepG2 cells (32), mice (33, 34), and pigs (35) revealed that loss of *ASGR1* increases the levels of surface LDLR through both PCSK9-independent (32) and dependent (33–35) mechanisms. Data from cultured HepG2 cells suggest that ASGPR directly binds LDLR, facilitating LDLR degradation; consequently, loss of *ASGR1* led to elevated numbers of LDLR on the cell surface (32). Alternatively, in mouse, *Asgr1* was found to release sterol regulatory element–binding proteins (SREBPs) from the ER (33). Thus, loss of *Asgr1* led to ER-trapped SREBP, which in turn decreased levels of PCSK9 and increased surface LDLR numbers (33). In contrast, a separate mouse study indicated that loss of *Asgr1* does not affect LDLR but instead prevents the degradation of the liver X receptor (LXR). The increased levels of LXR lead to the upregulation of the ATP-binding cassette (ABC) transporters *Abca1* and *Abcg5/g8*. *Abca1* facilitates reverse cholesterol transport to HDL, while *Abcg5/g8* shuttle cholesterol towards bile synthesis and fecal secretion (36, 37), thus their upregulation results in reduced cholesterol in the liver (34). Both studies reported that loss of *Asgr1* led to lower serum cholesterol and triglycerides, but the effects on lipoproteins contradicted each other (33, 34). In contrast to these more recent studies, no effect on lipoprotein levels was described in the first mutant mouse model of *Asgr1* (38). The differences noted between these different studies may have been due to the specific mutations present in the different mouse models, suggesting that further comparative analyses are warranted.

While these studies were ongoing, we independently investigated the influences of *ASGR1* on lipoprotein metabolism in zebrafish. Zebrafish are an underutilized, but powerful, model organism to study lipoprotein metabolism (39, 40). In contrast to mice, which primarily transport lipids through the high-density lipoproteins (41), zebrafish show a similar lipoprotein profile to humans in that they transport a bulk of their lipids through B-lps and express the cetp gene (42). Some groups chose the pig as their model organism as its lipoprotein profile is also more similar to human than rodents; however, mechanistic studies are difficult in a large, slow-developing organism (35). In contrast, cell culture-based studies are excellent for mechanistic details; however, they lack the context of a full organism and the circulatory system. Many processes, especially during development, can be examined directly since zebrafish reproduce externally in high numbers, with larvae that are optically transparent and develop rapidly. Further, critical genes involved in mammalian lipoprotein metabolism have been identified in zebrafish (43–49), and the availability of customizable diets (50, 51) and a reporter to measure lipoproteins in zebrafish (52) make zebrafish a suitable model organism for metabolic diseases (reviewed by 39, 40, 53-57).

Here, we identified the zebrafish ortholog for *ASGR1*, *asgr1a,* and generated two knockout lines using CRISPR/Cas9. Adult zebrafish *asgr1a* mutants challenged with a Western Diet (WD) developed less steatosis and stored less triglyceride in the liver while showing increased cholesterol levels in their feces compared to control animals. Our results highlight the importance of ASGR1 in whole body lipid.

## Results

### In silico analysis suggests complicated evolution of asgr1 in zebrafish

While zebrafish share 70 % of the human coding genes (58), a BLAST search of the human ASGR1 amino acid sequence using standard default settings returned mostly unannotated genes (9 out of 20, Fig. 1 A) and several genes of the C-type lectin family (6 out of 20), but no immediately obvious ASGR ortholog. To identify the zebrafish ortholog for ASGR1, we narrowed down potential candidate genes using various criteria. First, we selected the top three results of non-annotated genes, *si:ch73-361h17.1*, *si:ch211-283g2.2*, and *si:ch211- 283g2.3,* which all showed similar identity scores (36.63 - 40.60 %). We also selected the unannotated gene *BX640512.3,* which was predicted to belong to the C-type lectin-binding protein family by the genome browser (25.11 % identity). Third, we included a recently described gene zebrafish hepatic lectin *si:ch211-170d8.5* (32.64 % identity) (59). *ASGR1* shares considerable similarities with *ASGR2* and *CLEC10A* both in function and sequence (5); thus, we used the pair-wise alignment of all zebrafish candidates using ClustalW, revealing that *si:ch73-361h17.1* had the highest overall similarity with ASGR1 (42.4 %) (Supp. Fig. 1).

**Figure 1:**
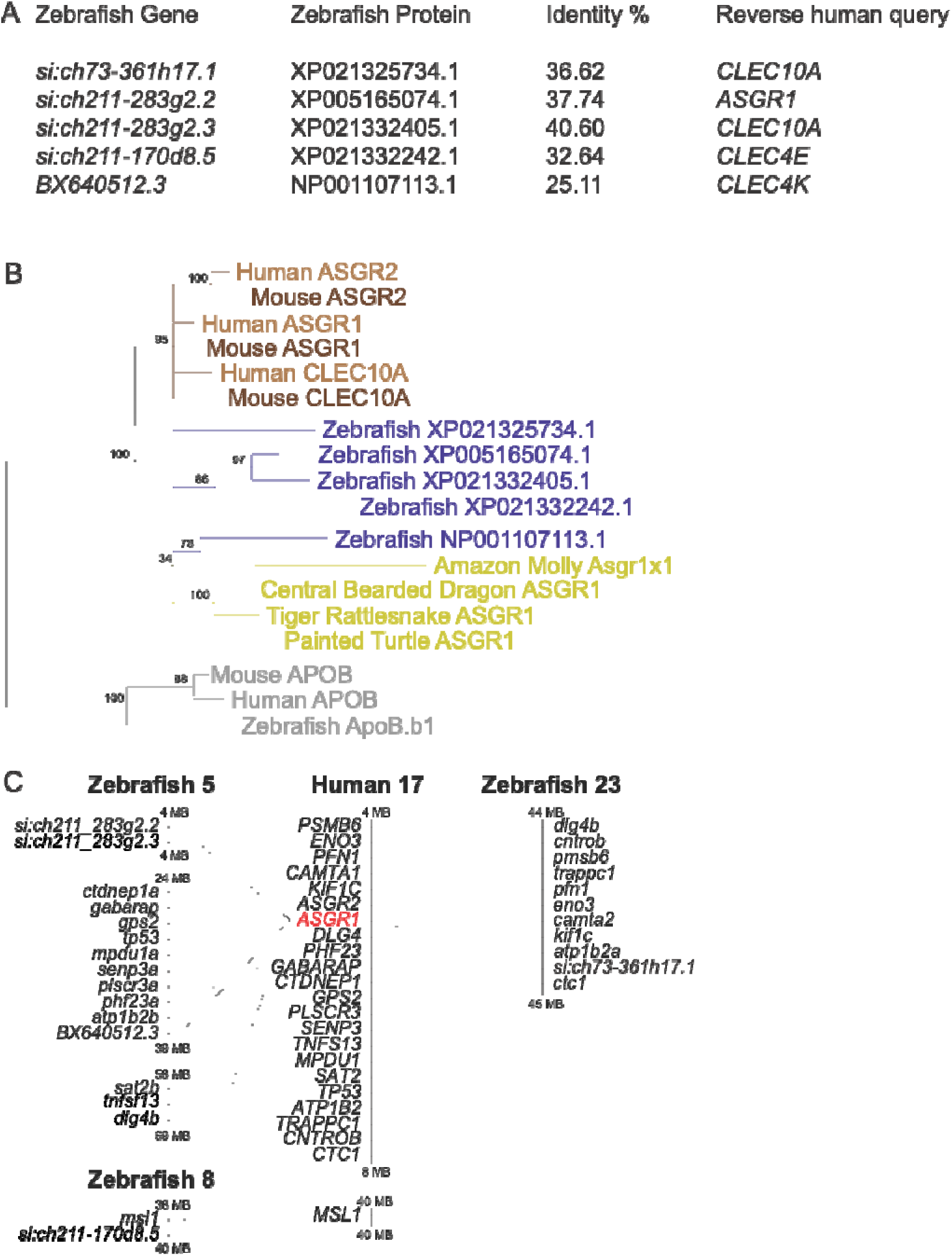
Gene origin analysis for zebrafish asgr1 paralogs. (A) BLAST of the human ASGR1 protein sequence to zebrafish genome using standard settings in ensemble.org shows low similarity. Reverse BLAST is inconclusive. (B) Phylogenetic tree based on amino acid sequences of human and mouse ASGR1, ASGR2, and CLEC10A to possible zebrafish orthologs and other non-mammalian ASGR1 orthologs. APOB sequence serving as outgroup. (C) Syntenic analysis of human Chromosome 17 and possible zebrafish orthologs, not to scale. Light grey lines show synteny, black lines connect zebrafish asgr1 candidates.

We turned to phylogenetic analysis based on amino acid sequences to further narrow down the five potential candidate genes. Human, mouse, and zebrafish APOB were chosen as an outgroup to help the MAFFT multiple alignment tool (60) determine relations within the ASGR1 sequences. In addition to ASGR1, we again included ASGR2 and CLEC10A from both the mouse and human in our analysis (5). Based on the multiple alignments, MAFFT generated a neighborhood-joining tree with bootstrap sampling. The mouse and human orthologs of ASGR1, ASGR2, and CLEC10A all form a clade with high bootstrap values, indicating that these mammalian orthologs share a common ancestor that probably diverged at some time after the evolutionary division between ray-finned and lobe-finned fishes (Fig. 1 B).

Since the zebrafish genome underwent a duplication event (61, 62), a zebrafish gene having several orthologs to a human gene is not uncommon. Syntenic analysis examines one gene and its neighboring genes, as they tend to remain in proximity throughout evolution (63). Therefore, we identified the genes located near the *asgr1* candidate genes. *ASGR1* is located on human chromosome 17 at 7.17 MB. Syntenic analysis showed that the genes surrounding human *ASGR1* (4 to 40 MB examined) could be found in blocks on the zebrafish chromosomes 5 (*si:ch211-283g2.2*, *si:ch211-283g2.3*, and *BX640512.3)* and 23 (*si:ch73-361h17.1)*. The candidate gene *si:ch211-170d8.5* is located on chromosome 8, which has minor similarities to the human chromosome 17 (Fig. 1 C). We conclude from this analysis that the candidate genes on chromosomes 5 and 23 result from the genome duplication and the genes on chromosome 5 further underwent a tandem duplication, which is common in zebrafish (64).

### mRNA expression pattern of candidate zebrafish asgr1 paralogs

In addition to sequence and syntenic analyses, the closest zebrafish ortholog to a human gene is often identified by examining the mRNA expression pattern with whole-mount *in situ* hybridization (WISH). Determination of mRNA expression patterns is crucial when identifying zebrafish orthologs since the tetrapod genome duplication often leads to silent copies of genes in zebrafish (61, 62, 65–70). We performed WISH with specific anti-sense and sense riboprobes against all candidate mRNA sequences on whole zebrafish larvae. Our goal was to identify the candidate gene with liver-specific expression since ASGR1 shows liver-specific expression in humans while CLEC10A is ubiquitously expressed (5). Only the probes for *si:ch73-361h12.1* and *si:ch211-170d8.5* showed liver expression. *Si:ch73-361h12.1* signal was detected in the liver at 3 dpf (n = 29 of n_total_ = 31), while only a subset of larvae showed *si:ch211-170d8.5* expression at 3 dpf (n = 3 of n_total_ = 27). At 5 dpf, both probes exhibited strong liver-specific mRNA expression (Fig. 2). Sense controls for all riboprobes showed no signal (Suppl. Fig. 2).

**Figure 2:**
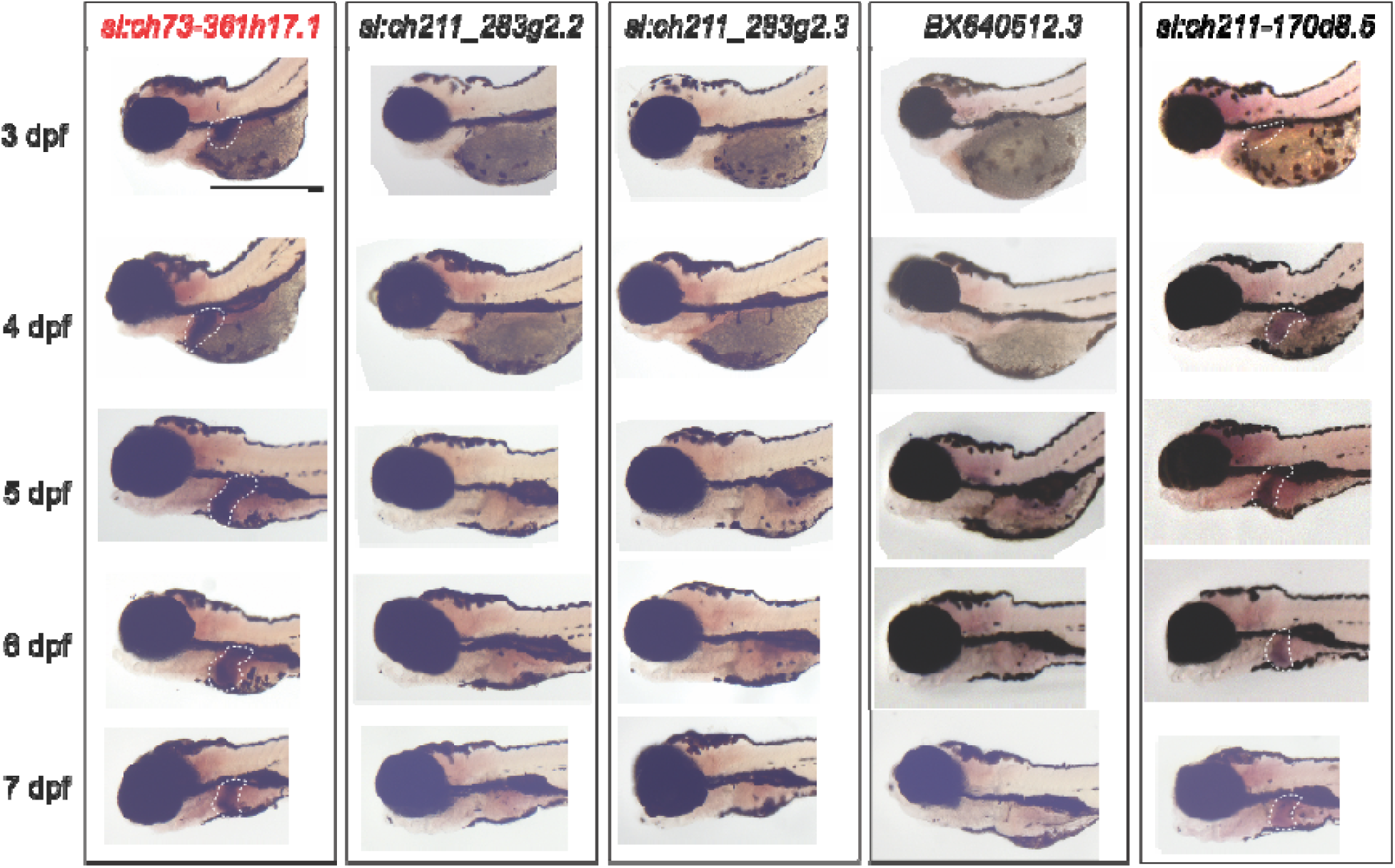
mRNA expression of *asgr1* candidate genes during larval stages. Whole-mount in situ hybridization (WISH) for riboprobes of *si:ch73-361h17.1* and *si:ch211-170d8.5* shows signal during 3 - 7 dpf. The signal is confined to the developing liver (white dotted outline). For *si:ch211_283g2.2*, *si:ch211_283g2.3*, and *BX640512.3* no signal was ever detected. WISH was performed three independent times on probes for all candidate orthologs at all developmental stages, n ≥ 6 in each experiment. *Si:ch73-361h12.1* showed liver signal consistent 36/36 every experiment starting at 4 dpf and 29/31 at 3 dpf. *Si:ch211-170d8.5* showed consistent signal 22/29. starting 4 dpf and 3/27 at 3 dpf. Scale bar 1mm.

These data support *si:ch73-361h12.1* being the ASGR1 ortholog, given its higher similarity to human ASGR1, its syntenic position, and gene expression. The correct nomenclature of these genes was determined in conjunction with the Zebrafish Model Organism Database (ZFIN). The active gene is *si:ch73-361h12.1,* named *asgr1a,* and the non-expressing duplicate is *BX640512.3,* named *asgr1b.* The paralogs *si:ch211-283g2.2* and *si:ch211-283g2.3,* which do not actively express mRNA, most likely evolved by tandem duplication and are named *asgr1c.1* and *asgr1c.2*, respectively. *Si:ch211-170d8.5* does show liver-specific expression; however, its distance in both phylogenic and syntenic analysis supports its naming as *asgr1-like* (*asgr1l*).

### Generation of two asgr1a mutant zebrafish lines

Having identified the likely zebrafish ASGR1 ortholog, we generated two independent *asgr1a* mutants using CRISPR/Cas9. For the first mutation, we targeted the start codon of *asgr1a* (Fig 3 A). This produced a 5 base pair (bp) deletion in exon 2 (*asgr1a15_20del*) with a predicted premature stop codon after 26 amino acids. This allele is now designated as Carnegie c830 (*asgr1a^c830/c830^*) (Fig. 3 B-D). Sequencing of gDNA and cDNA (by RNAseq analysis of the mutated locus) confirmed the deletion (Supp Fig. 3 A). Genotyping this small deletion mutation is challenging; however, heterozygous animals can be identified readily by heteroduplex formation (71, 72) (Fig. 3 B). In addition to the challenges of genotyping *asgr1a^c830/c830^*, we were concerned that alternative start sites could mask a phenotype since several potential start codons could lead to a partially functional protein (73–75). A second independent mutation in the predicted C-type lectin binding domain was generated to address this concern. This domain is essential for binding glycoproteins; hence, its amino acid sequence shows more conservation than the other functional domains (Supp. Fig 1 B). By injecting three CRISPR guides spaced several base pairs apart, a 33 bp mutation was introduced in exon 9 (*asgr1a791_824del*), which was assigned Carnegie c865. This mutation, *asgr1a^c865/c865^*, leads to the loss of nine conserved amino acids and changes one conserved amino acid (Fig. 3 C, Supp Fig 3 B), shortening the full-length protein to 288 amino acids long compared to the wild-type sequence of 299 amino acids (Fig. 3 B-D). It is known that silent paralogs can turn on expression to compensate for losing the usually active gene (75, 76).

**Figure 3:**
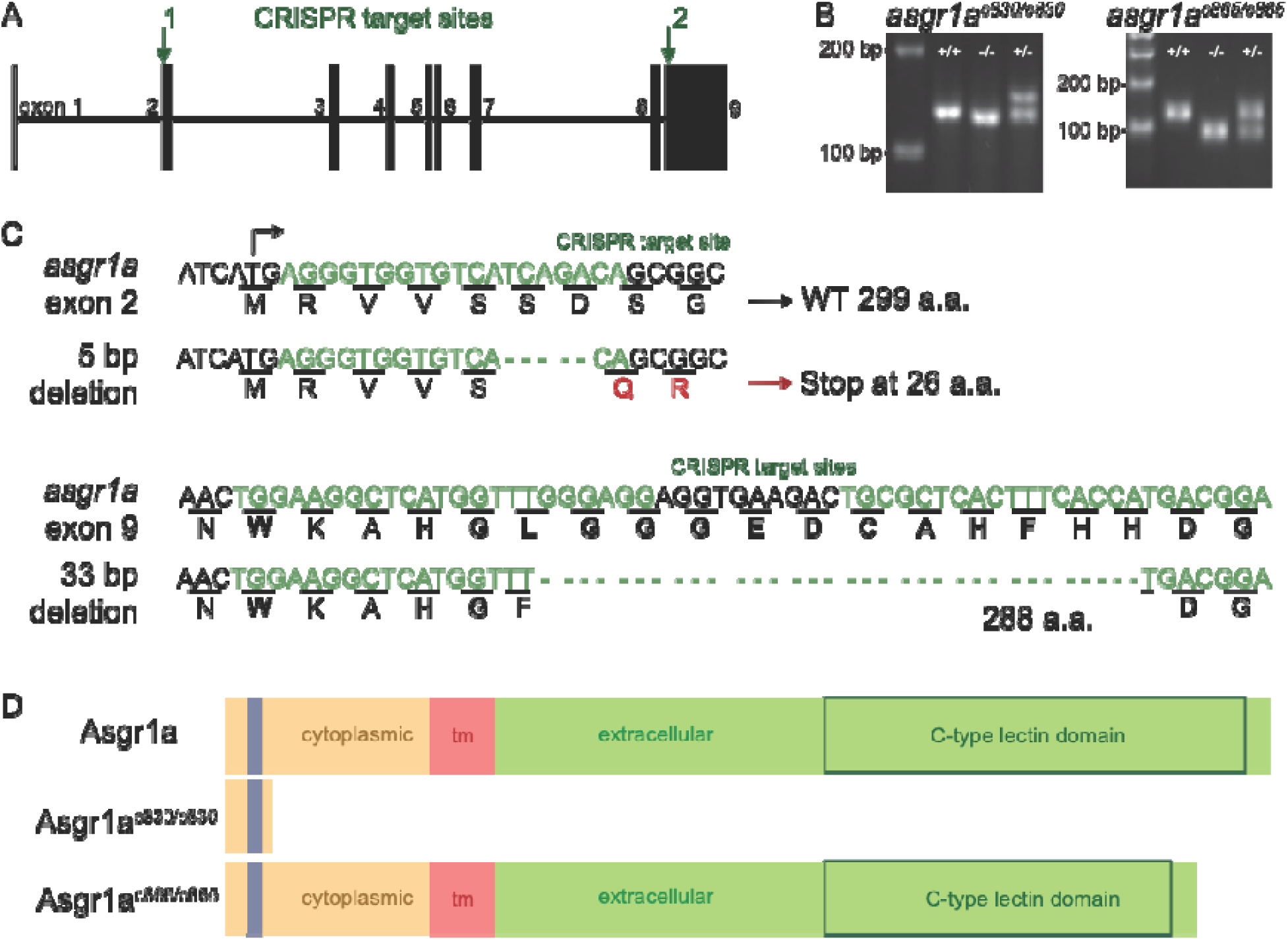
Generating two mutations in *asgr1a* using CRISPR/Cas9. (A) Schematic of *asgr1a* gene showing the CRISPR target sites in exon 2 and 9. (B) Representative images of genotyping assays for both mutations showing heteroduplex formation in *asgr1^830/+^*. (C) Detailed view of *asgr1a^c830/c830^*(5 bp deletion in exon 2) and *asgr1a^c865/c865^* (33 bp deletion in exon 9). Conserved amino acids to human ASGR1 are highlighted in bold. Red indicates the change in amino acid sequence, green letters denote the sequence of CRISPR target sites. Abbreviations: (bp) base pair, (a.a.) amino acid. (D) Predicted protein length and domains in WT and both *asgr1a* mutations. Orange for the cytoplasmic region with the blue endocytosis motif, red for the transmembrane domain, green for the extracellular domain of the protein with a box highlighting the predicted area of the C-type lectin domain. Assignments were performed based on pairwise alignment of the amino acid sequence to human ASGR1 and its known functional domains.

### Asgr1a mutants secrete excess dietary lipids

The LipoGlo reporter system allows for the quantification of B-lp numbers and estimation of B-lp sizes in zebrafish (52). The reporter was designed to examine B-lps in individual zebrafish larvae (52, 77) by expressing nano-luciferase (NanoLuc) fused to the endogenous C-terminus of ApoBb.1, the zebrafish ortholog for human APOB. Each B-lp requires a single copy of APOB as a structural protein for proper function (78), which is not interchangeable. Therefore, the relative luminescence signal from the NanoLuc provides a one-to-one read-out for the total amounts of B-lps (52). To measure the impact of *asgr1a* loss on B-lp metabolism in zebrafish using the LipoGlo Counting assay, *asgr1a^c830/c830^*and *asgr1a^c865/c865^* were crossed into the LipoGlo reporter line. No detectable change in B-lp number or particle size was detected in the *asgr1a* mutants and their wild-type siblings during larval and juvenile stages (Supp. Fig. 4).

Homozygous mutants for both *asgr1a* mutations are viable, and no changes in mass or length at six months of age were observed when fed a normal diet (Supp. Fig 5 A, B). Recent mouse studies reported that the loss of *Asgr1* led to less hepatic steatosis due to excess dietary cholesterol secreted in the feces (33, 34). Thus, adult zebrafish were challenged with a WD for 6 days. *Asgr1a^c830/c830^*, *asgr1a^c865/c865^* mutant fish, and WT controls were fed a custom WD provided by Sparos. After the morning feed on the sixth day, fish were separated, and feces were collected before the fish were dissected to obtain plasma and livers for analysis.

Females are not desired when studying hepatic phenotypes as their hepatic structure is highly variable and dependent on the timing of their hormonal cycle (78). Thus, unsurprisingly, females showed no phenotype in response to the WD feeding (Supp Fig. 5 C-H). The LipoGlo reporter system can be adapted to measure B-lp levels in 1 µL of adult zebrafish plasma (51). Neither *asgr1a* mutation affected B-lp levels or particle sizes in male zebrafish measured with the LipoGlo-Counting and -Electrophoresis assays (Fig. 4 A-C). To examine the levels of liver steatosis, pieces of liver were analyzed by Hematoxylin and eosin (H & E) and High-Pressure

**Figure 4:**
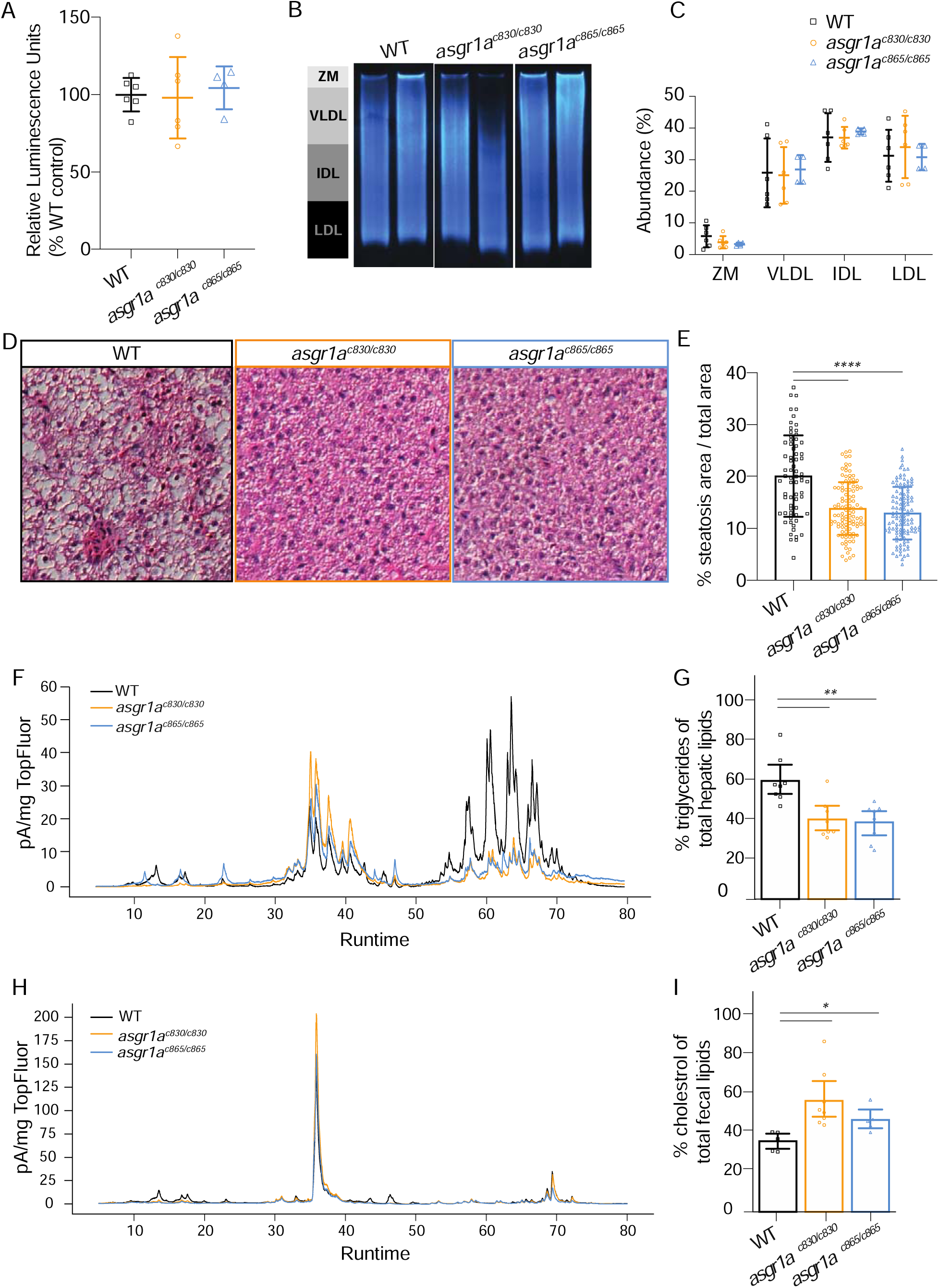
Asgr1a mutants show less hepatic steatosis and lower levels of hepatic triglycerides but excrete more cholesterol in response to excess dietary cholesterol. Male zebrafish received a WD for 6 days before blood, liver, and feces were collected. (A) LipoGlo-Counting of plasma samples did not show any changes in B-lp quantity between asgr1a mutants and control animals. Two independent experiments, n = 2 – 3 fish each; two-way ANOVA followed by Tukey’s multiple comparison test. Experiments were normalized to their control group and shown in % to the control group. (B) Representative images of LipoGlo-Electrophoresis on plasma B-lps. Zero mobility (ZM) and B-lp classes (VLDL, IDL, LDL) can be distinguished and analyzed. (C) Quantification of LipoGlo-Electrophoresis assay showing no changes in B-lp size distribution. Two independent experiments, n = 2 - 4 fish/genotype; two-way ANOVA followed by Tukey’s multiple comparison test. (D) Representative H & E images of liver steatosis. (E) asgr1ac830/c830 (13.82 % ±5.079) and asgr1ac865/c865 (12.90 % ± 5.058) mutants develop significantly less hepatic steatosis compared to control animals (20.06 % ± 7.829). n = 9 from three independent experiments; two-way ANOVA followed by Tukey’s multiple comparison test. (F) Example trace of one sample of an asgr1a mutant (blue and orange) and control (black) livers. (G) Peak analysis of three independent experiments, n = 9, revealed less hepatic triglycerides in asgr1ac830/c830 (40% ±9.7) and asgr1ac865/c865 (38 % ±9.3) mutants compared to controls (60 % ±11). (H). Example trace of one sample of an asgr1a mutant (blue and orange) and control (black) fecal matter. (I) Peak analysis of three independent experiments, n = 9, revealed more fecal cholesterol in asgr1ac830/c830 (67% ±17) and asgr1ac865/c865 (56 % ±7.4) mutants compared to controls (43 % ±5.8). Statistical analysis as follows (E) Analyzed using two-way ANOVA followed by Tukey’s multiple comparison test. (G and I) Significance was computed using two independent t-tests with the Bonferoni correction. * = p < 0.05; ** = p < 0.01; **** = p < 0.0001

Liquid Chromatography (HPLC). H & E analysis showed that there was a significant reduction in hepatic steatosis in males of both mutants for *asgr1a* compared to the controls in three independent experiments (Fig. 4 D, E, Supp Fig. 6; control: 20.06 % ± 7.83; *asgr1a^c830/c830^*: 13.82 % ± 5.08; *asgr1a^c865/c865:^*12.90 % ± 5.06). HPLC analysis of the same liver samples revealed a decrease in triglycerides in both *asgr1a* mutants by about 20 % (Fig. 4 F, G control 60 % ± 11; *asgr1a^c830/c830^*: 40 % ± 9.7; *asgr1a^c865/c865:^* 38 % ± 9.3). Feces collected from the same fish and analyzed by HPLC revealed that *asgr1a* mutants excreted more cholesterol than the control animals (Fig. 4 F, G control 43 % ± 5.8; *asgr1a^c830/c830^*: 67 % ± 17; *asgr1a^c865/c865:^* 56 % ± 7.4). Though we performed the same analyses in females, we found no phenotype in response to the WD feeding (Supp Fig. 5 C-G), possibly due to the metabolic changes that occur with their 5-day-hormonal cycle (79, 80).

## Discussion

Concurrent with this study, four groups started investigating the loss of ASGR1 after it was reported to reduce LDL-cholesterol and the development of cardiovascular disease (31). Experiments with cultured cells and in pigs indicated that lowered lipoprotein levels in *ASGR1* mutants were achieved by increased surface LDLR in hepatic cells (32, 35). In mice, two complete knockouts for *Asgr1* were reported by two research groups. One mouse study indicated that *Asgr1* acts on the LDLR through the SREBP pathway (33). The second study performed in mice reported a mechanism via LXR and did not see an effect on LDLR (34). While these two studies show contradicting results about total lipoprotein numbers and profiles and thus raise more questions, both reported an increase in fecal cholesterol in *Asgr1* mutants.

We used a combination of phylogenetic and syntenic analysis, followed by *in situ* hybridization, to detect the active ortholog of *ASGR1* in zebrafish. These methods are often used together to help find functional, active zebrafish orthologs of human genes since zebrafish often have several ohnologs of genes due to genome duplication and random tandem duplication (61, 62, 65–70). While we identified the zebrafish orthologs for *ASGR1*, simultaneously, *asgrl1* (59) and *asgr1a* (81) were described by the Li lab at the Ocean University of China. In addition to expression pattern analysis by WISH and qPCR, both genes were described to be involved in the immune response and clearance of pathogens from circulation (59, 81), which are known functions of the human ASGR1 (14–17).

We generated two independent mutations in *asgr1a* and did not see changes in lipoprotein numbers or particle sizes. This is contradictory to the GWAS findings that heterozygous loss of *ASGR1* in humans is correlated with reduced plasma LDL levels (31). As we could not detect any phenotype of *asgr1a* mutants under normal conditions (Supp. Fig 4, 5 A, B), we challenged the metabolism of zebrafish by feeding the zebrafish a custom WD. Indeed, when feeding the zebrafish a WD for 6 days, *asgr1a* mutants developed less steatosis and instead excreted the additional dietary lipids, especially cholesterol, in their feces, similar to some mouse phenotypes (33, 34).

ASGRP is a receptor for many different particles, not just lipoproteins. Thus, part of the difficulty when establishing a mechanism of action for *asgr1a* on hepatic lipids is the polyvalent effect of ASGR1 on many different pathways. A recently published drug-target Mendelian randomization study examined the effects of genetically mimicked ASGR1 inhibitors. Concurrent with our results that loss of ASGR1 affects a multitude of pathways, they found reduced levels of TG and APOB and also changes in insulin-like growth factor 1, albumin, and calcium (82). These side effects are not surprising, given the polyvalence of genes that we saw affected by the *asgr1a* mutation. In summary, our results in zebrafish emphasize importance of ASGR1 in modulating total fecal sterol levels, and hepatic steatosis may underlie human GWAS data indicating that attenuated ASGR1 function is cardioprotective. However, the mechanistic underpinnings that can explain the relationship between ASGR1 function and fecal sterol levels remain to be described.

## Methods and Materials

### Gene ancestry analysis

Sequence, syntenic, and phylogenetic analyses were performed to identify the asialoglycoprotein receptor 1 (asgr1) ortholog in zebrafish. All genetic and amino acid sequences were obtained from the National Center for Biotechnology Information (https://www.ncbi.nlm.nih.gov) and the Ensembl Genome Browser (ensembl.org).

Possible zebrafish orthologs were identified by querying the human protein sequence of *ASGR1*, transcript 201, on ensemble.org using BLASTp (standard settings: “Search Against”: Protein database “Proteins(Ensembl)”; “Search Sensitivities” set to “normal”, no Additional configurations) to the zebrafish RefSeq protein database (83, 84). The top three BLAST hits were recorded. A fourth candidate gene was chosen by looking at C-type lectin-containing protein families. The fifth candidate was a zebrafish hepatic lectin recently described by (59). Retro-BLASTp for all five potential zebrafish orthologs was performed. The amino acid sequences of human, mouse, zebrafish, central bearded dragon, western painted turtle, amazon molly, and tiger rattlesnake for ASGR1, and the human and mouse orthologs for ASGR2 and CLEC10A were aligned using the iterative MAFFT multiple sequence alignment tool (85, 86) to examine the phylogenetic evolution. For accurate guide tree building, ApoB protein sequences for human, mouse, and zebrafish were chosen as outgroups (60). The G- INS-1 strategy was selected using standard settings (“Try to align gappy regions anyway”; Parameters: Scoring matrix for amino acid sequences: “BLOSUM62”; Gap opening penalty: “1.53”, Offset value: “0.0”; Score of N in nucleotide data: “(nzero) N has no effect on the alignment score”; Guide tree: “Default”; Mafft-homologs: “Use UniRef50 2019/Mar”), as it is recommended for accurate guide trees. Based on the alignment, the MAFFT tree server built a neighbor-joining tree with all gap-free sites, the JTT substitution model, and the estimation of site heterogeneity (alpha). Bootstrap resampling was set to 100, with bootstrap support ≥70 being interpreted as significant, and branches less than 50 were collapsed.

Optimal pairwise protein alignments between the human ASGR1 and all five potential zebrafish orthologs were determined with the ClustalW tool available in the MacVector software. BLOSUM was selected while other settings were kept standard. Identity and similarity scores for human ASGR1, ASGR2, and CLEC10A to all five potential orthologs were determined with multiple alignments, same settings as previously noted.

The syntenic relationship between the human ASGR1 and possible zebrafish gene loci was established using Ensembl Genome Browser by manually searching for zebrafish orthologs of genes located on human Chromosome 17, 4-40 MB (ASGR1 located at 7,173,431-7,179,370 MB).

### Zebrafish husbandry

Embryos were collected from the natural spawning of group crosses with two females and two males. Embryos were staged (87) and kept in embryo medium at 28 °C on a 14:10 light: dark cycle until 6 days post fertilization (dpf). Larvae were then moved to the fish facility (14:10 light: dark cycle) and fed with GEMMA Micro 75 (Skretting) three times a day until 14 dpf, GEMMA Micro 150 three times a day, and Artemia once daily from 15 - 42 dpf. Adult zebrafish were fed once daily with ∼3.5% body weight Gemma Micro 500 (Skretting USA). Fish challenged with a Western Diet were fed as before, but with Sparos custom Western Diet. Fish were fed with the WD for 6 days before downstream experiments.

The Carnegie Institution Department of Embryology Animal Care and Use Committee (Protocol #139) approved all animal experiments.

### Whole-mount in situ hybridization

Whole-mount *in situ* hybridization (WISH) was performed as previously described (88) to identify the expression of ortholog candidate genes for asgr1 in zebrafish. Unique sequences for *asgr1a*, *asgr1b*, *asgr1c.1*, and *asgr1c.2* were obtained from Genewiz (sequences listed in supplementary material). The riboprobe for *asgrl1* was generated from cDNA according to (59). All riboprobes were cloned using TOPO® cloning into pCRII (Thermo Fisher Scientific, K461020). Translation with Sp6 and T7 (Millipore Sigma, 10810274001 and 10881767001) generated sense and antisense digoxigenin-labeled riboprobes (Anti-Digoxigenin-AP, Fab fragments, Roche 11093274910). WISH was performed three times with an n > 5 on AB larvae stages 3 - 7 dpf and on 6 dpf larvae carrying either the *asgr1a^c830/c830^* or the *asgr1a^c865/c865^* mutation. Fish were imaged using a Nikon SMZ1500 microscope with HR Plan Apo 1x WD 54 objective, Infinity 3 Lumenera camera and Infinity Analyze 6.5 software.

### CRISPR/Cas9 mediated knockout

CRISPR target sites were identified with the help of the UCSC Genome Bioinformatics tool (https://genome.ucsc.edu/). The zebrafish genome was chosen, the locus for *asgr1a* was entered, and “CRISPR” under “Genes and Predictions” was set to “show”. Potent PAM sites were selected, and by clicking on the site, the guide sequence was displayed. Areas with high efficiency were chosen and ordered from Eurofins Genomics (supplementary primers 7 – 10). Oligos were resuspended into a 10 µM working stock and set up in a PCR reaction to prime to the CRISPR tail (primer 13) with dNTPs and high-fidelity polymerase (Phusion polymerase, Thermo Scientific #F-530S) (T_a_ = 55 °C, extension time 30 s, 15 cycles). After purification using the Zymo DNA Clean and Concentrate kit (Zymo Research, D4013), *in vitro* transcriptions into sgRNA were performed using the MEGAshortscript T7 kit (Thermo Fisher Scientific, AM1354). The sgRNAs were cleaned using the Zymo RNA clean and concentrate (Zymo Research, R1013). AB zebrafish were injected with a 2 nL injection mix (1500 ng/µL) at the 1-cell stage and raised to adulthood for founder identification. By genotyping their offspring, founders were identified.

### DNA extraction and genotyping

Genomic DNA was obtained by adapting the HotSHOT DNA extraction protocol (89). Adult fin clips were heated to 95 °C for 20 min in 50 µL of 50mM NaOH, single larva in 20 µL. After cooling to room temperature, 1/10^th^ volume of 1 M Tris pH 8.0 was added.

Genotyping primers for both *asgr1a* mutants were designed using ApE and synthesized by Eurofin Genomics. The locus around the 5 bp mutation in exon 2 of *asgr1a* was amplified using primers 1 and 2 at 0.5 µM concentration (T_a_ = 56 °C, extension time 20 s, 36 cycles). The WT band is 130 bp, and the homozygous mutant band is 125 bp. Heterozygous mutants show heteroduplex behavior in gel electrophoresis: bands run at approximately 128 bp and 140 bp. The locus of the 33 bp mutation in exon 9 of *asgr1a* was amplified using primers 3 and 4 (0.5 µM primer, T_a_ = 52 °C, extension time 40 s, 34 cycles). The WT band is 129 bp, and the homozygous mutant band is 96 bp. Genotyping of ldlr mutants was performed using primers 5 and 6 (T_a_ = 64 °C, extension time 30 s, 36 cycles). The WT band is 128 bp, and the homozygous mutant band is 118 bp. For the ApoBb.1-NanoLuc genotyping protocol, see (52).

### LipoGlo assays

See (52) for detailed LipoGlo methods. All reagents were obtained from Promega Corp., (N1110; (90). Larval, juvenile, and adult *asgr1a^c830^* mutants, as well as adult *asgr1a^c865^* mutants carried one copy of the LipoGlo reporter (*apoBb.1^Nluc/+^*), while larval experiments with *asgr1a^c865^* animals carried two copies (*apoBb.1^Nluc/Nluc^*).

To measure lipoprotein levels in adults, blood was collected at the time of dissection by cutting of the fin and squeezing the body to collect the blood using EDTA-coated Kunststoff Kapillaren tubes (Sanguis Counting, Nümbrecht, Germany). The blood was centrifuged in a 1.5 mL Eppendorf tube for 5 min at 5000 g at 4 °C, and the plasma (1-2 µL) was pipetted into a new 1.5 mL Eppendorf tube and snap-frozen on dry ice. For larval LipoGlo assays, larvae were dispensed into 96-well plates (USA Scientific, #1402-9589) for homogenization in a total volume of 100 µL of 2x B-lp stabilization buffer (40 mM EGTA, pH 8.0, 20% sucrose + cOmplete Mini, EDTA-free protease inhibitor (Sigma, 11836170001)). Juveniles were dispensed into 96-well opaque white OptiPlate (Perkin-Elmer, 6005290) in a total volume of 200 µL of B-lp stabilization buffer. Homogenate was generated by sonication in a microplate-horn sonicator (Qsonica Q700 sonicator with a Misonix CL-334 microplate horn assembly) and kept on ice for immediate use before being stored at -20 °C. LipoGlo-Counting levels were obtained by mixing 1 µL of frozen plasma with 99 µL of diluted 1x B-lp stabilization buffer or by mixing 40 µL of larval or juvenile homogenate with 40 µL of diluted LipoGlo buffer (1:3 NanoGlo buffer: PBS + 0.5 % NanoLuc substrate) in a 96-well opaque black OptiPlate (Perkin-Elmer, 6005270).

All plates were read within 5 min of buffer addition. The plate for *asgr1a^c830/c830^* was read on a SpectraMax M5 plate reader (Molecular Devices) set to top-read chemiluminescent detection with a 500 ms integration time. *Asgr1a^c865^* LipoGlo-Counting plates and adult plasma LipoGlo-Counting samples were read on the BioTek Synergy H1 (Agilent, Santa Clara, CA) set to Luminescence Endpoint, 500 ms integration time.

Adult plasma LipoGlo-Electrophoresis reads vary drastically between experiments. To normalize for this, each experiment was normalized to its own control group by calculating the % of each value to the mean of the control.

LipoGlo-Electrophoresis was performed according to (52). Native-PAGE were imaged using the Odyssey Fc (LI-COR Biosciences) gel imaging system. For quantification, gels were analyzed using FIJI. Each lane was converted to a vertical plot profile. The Di-I LDL standard migration assigned areas for Zero Mobility (ZM), VLDL, IDL, and LDL.

### Tissue histology

Adult male zebrafish (12 mo age-matched for all genotypes) were fed a WD (custom product from Sparos, composition:) for 6 days. Fish were euthanized by high-dose exposure to tricaine (Sigma Aldrich, A5040-25). Through ventral sections, livers were removed as intact as possible and fixed in neutral-buffered formalin (Sigma, F8775) at RT for 24 hrs. The Johns Hopkins University Oncology Tissue Services performed sectioning and hematoxylin & eosin staining. Slides were imaged with a Nikon E800 microscope with 20×/1.4 Plan Apo Nikon objective and Canon EOS T3 camera using EOS Utility image acquisition software. Quantification of steatosis was performed using FIJI. Images were converted to RGB stacks, the threshold was set to 130 to 255, and the percentage area with steatosis was determined.

### HPLC

The quantification of lipid classes was performed by using high-performance liquid chromatography with a charged aerosol detector (HPLC-CAD). Tissues and fecal matter were suspended in 500 µL of 20 mM tris, 1 mM EDTA pH 7.8 lipid extraction buffer, and homogenized using a Fisher Scientific 550 Sonic Dismembrator. Using the Bligh-Dyer method (91), lipids were extracted from individual tissue homogenates and feces, then dried to 1% volume and resuspended in HPLC-grade isopropanol (>99.9% pure). The HPLC system (Thermo Fisher Scientific) with a C18 column and a CAD was used to analyze the lipid components of each sample. A gradient mobile phase was used for separating different lipid classes: 0 - 5 min = 0.8 mL/min in 98 % mobile phase A (methanol-water-acetic acid, 750:250:4) and 2 % mobile phase B (acetonitrile-acetic acid, 1000:4); 5 - 35 min = 0.8 - 1.0 mL/min, 98 - 30 % A, 2 - 65 % B, and 0 - 5 % mobile phase C (2-propanol); 35 - 45 min = 1.0 mL/min, 30 - 0 % A, 65 - 95 % B, and 5 % C; 45 - 73 min = 1.0 mL/min, 95 - 60 % B and 5 – 40 % C; and 73 - 80 min = 1.0 mL/min, 60 % B, and 40 % C. The amount of lipid in each sample was normalized to both the dry-weight of the samples and an internal lipid extraction control of TopFluor Cholesterol that was spiked into the lipid extraction solution and read on the fluorescence channel. For the figures generated in this paper, lipid classes were separated by runtime as established by (92), with fatty acids running from 5 - 30 min, phospholipids running from 30 - 45 min, and triglycerides and cholesteryl esters running from 45 - 75 min. Free cholesterol runs at 35 min, which can be seen in supplementary figure X. To calculate the quantity of each of these lipid classes, the area under the curve (AUC) was calculated in the given time windows after baseline correction established by running an isopropanol blank.

### Glucose Test

Animals of the following genotypes were fed a Western diet for 6 days before performing a glucose test: control: *apoBb.1^nluc/+^*; *asgr1a^c830/c830^*, *apoBb.1^nluc/+^*; *asgr1a^c865/c865^*, *apoBb.1^nluc/+^*. A control feeding with normal diet included *apoBb.1^nluc/+^*. All animals were age-matched to 6 months as best as possible. Forty-five minutes after the morning feed on day 6, animals were euthanized with tricaine and decapitated using a scalpel and cutting through the pectoral girdle. The cut severed the heart and the glucose strip was applied directly to the whole blood as described in (93). Measurements were taken with Alpha Trak 2 glucometer (amazon.com), set to Code 17, and Alpha Trak 2 test strips (amazon.com).

### Statistics

GraphPad Prism (GraphPad Software), or the Python packages Seaborn and StatAnnotate were used for graphing and statistical analysis. Outliers in all datasets were identified by ROUT and excluded from the analysis. Ordinary two-way ANOVA with Tukey’s multiple comparison test was performed on all developmental time courses of both *asgr1a* mutant LipoGlo experiments and the H & E quantification. Figure legends include sample sizes and details of statistical analysis.

