## Supplemental Figures for "Loss of *asgr1a* leads to the secretion of excess dietary cholesterol in zebrafish"

**S Figure 1: Protein similarity and identity scores of human ASGR1, ASGR2, CLECH10A to**
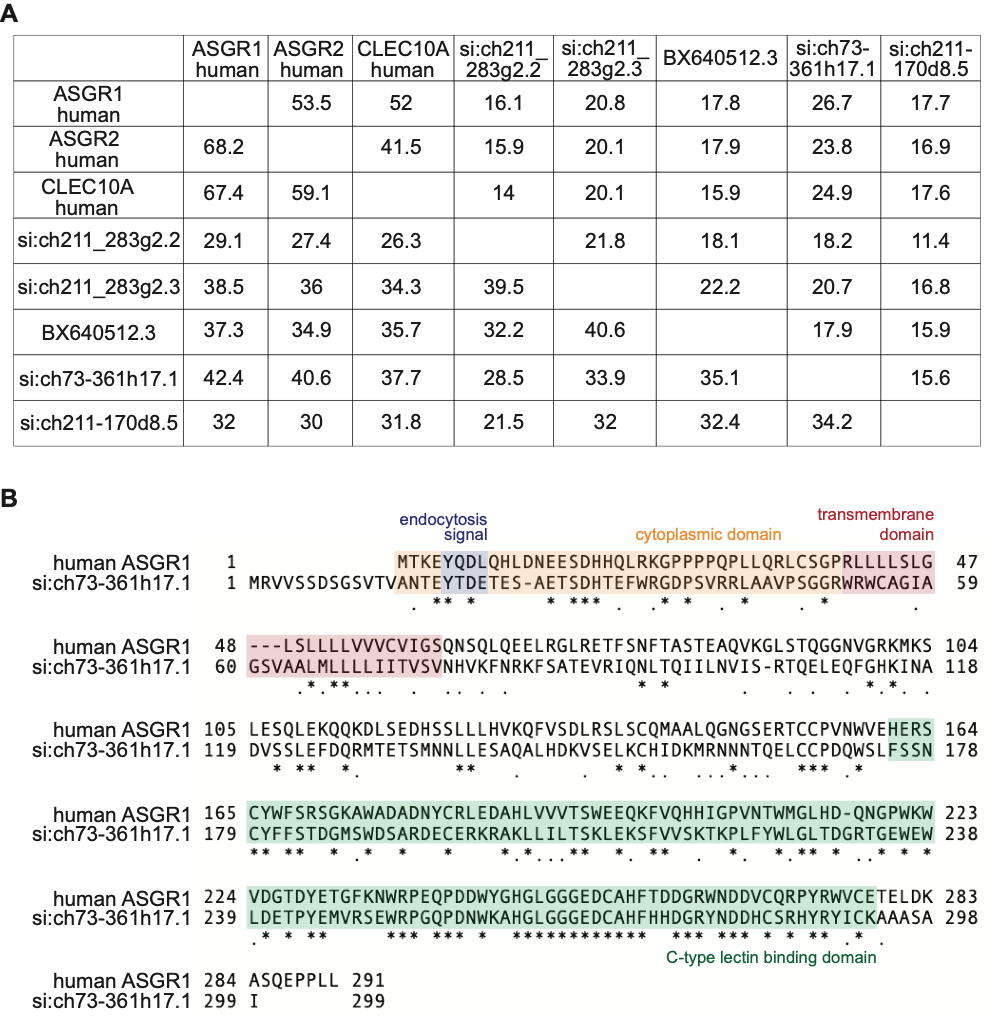
 **potential zebrafish orthologs.**

(A) Score matrix generated with MacVector by pair-wise aligning genes to each other based on ClustalW. Above diagonal identity scores are shown, below diagonal similarity scores. (B) ClustalW Alignment of human ASGR1 to si:ch73-361h17.1, functional domains marked with colors -orange: cytoplasmic domain, blue: endocytosis signal, red: transmembrane domain, green: C-type lectin binding domain. Identical amino acids marked by stars, functionally similar amino acids marked by dots.

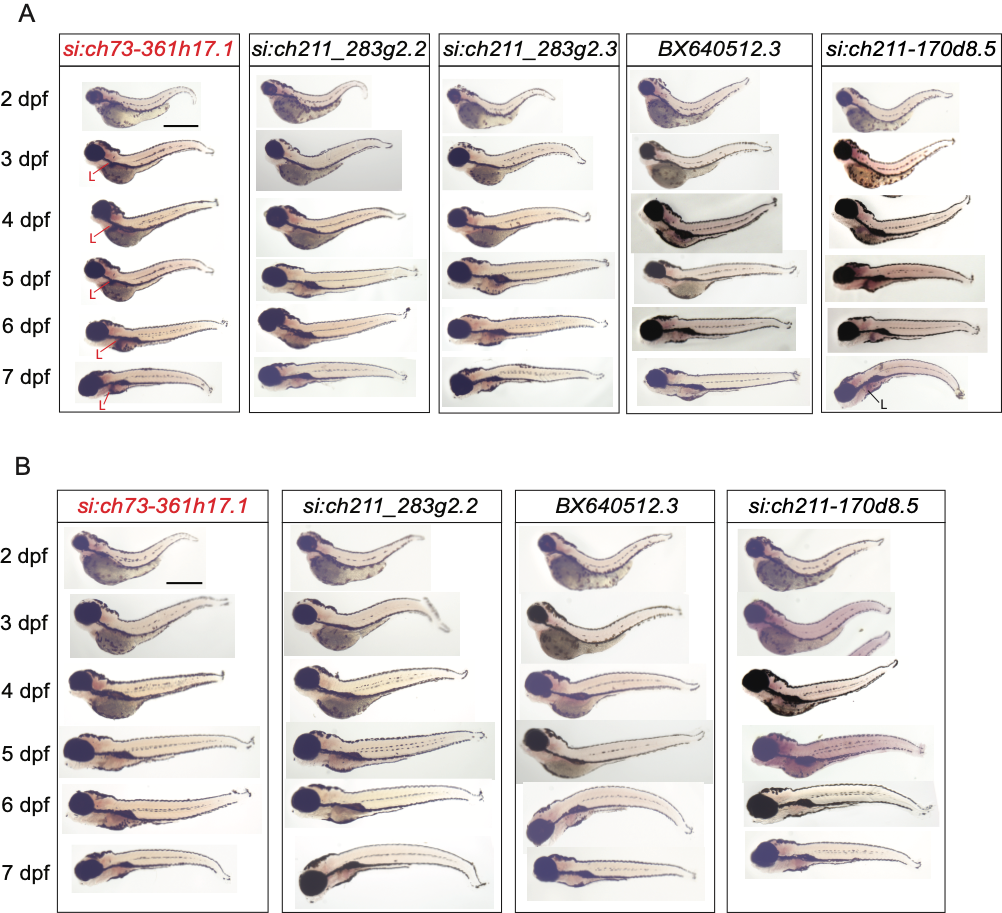
**S Figure 2: Whole larva mRNA expression of asgr1 candidate genes.**

(A) Full animal view of WISH for anti-sense riboprobes from Fig. 2. *Si:ch73-361h12.1* showed liver signal consistent 36/36 every experiment starting 4 dpf and 29/31 at 3 dpf, none at 2 dpf. *Si:ch211-170d8.5* showed consistent signal 22/29 starting 4 dpf and 3/27 at 3 dpf. (B) For each anti-sense probe, sense-probes were generated, except for BX45012.3. No signal could be detected in any larva when incubated with sense probes at 2 - 7 dpf. L: livers. Scale bar 1 mm.

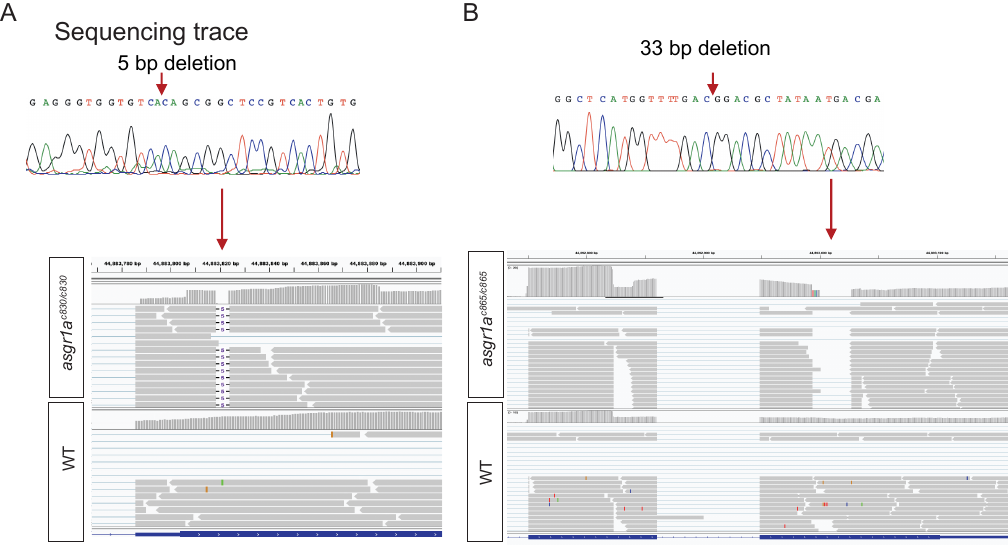

**S Figure 3: Confirmation of CRISPR/Cas9 induced deletion by genomic sequencing, RNAseq.**

(A, B) Trace file from sequencing genomic DNA obtained from single larvae mutant for (A) *asgr1a^c830/c830^* and (B) *asgr1a^c865/c865^*, confirming the deletions. Mapping RNAseq reads to the genome using IGV software, depicting the 5 bp and 33 bp deletion on RNA level, respectively.

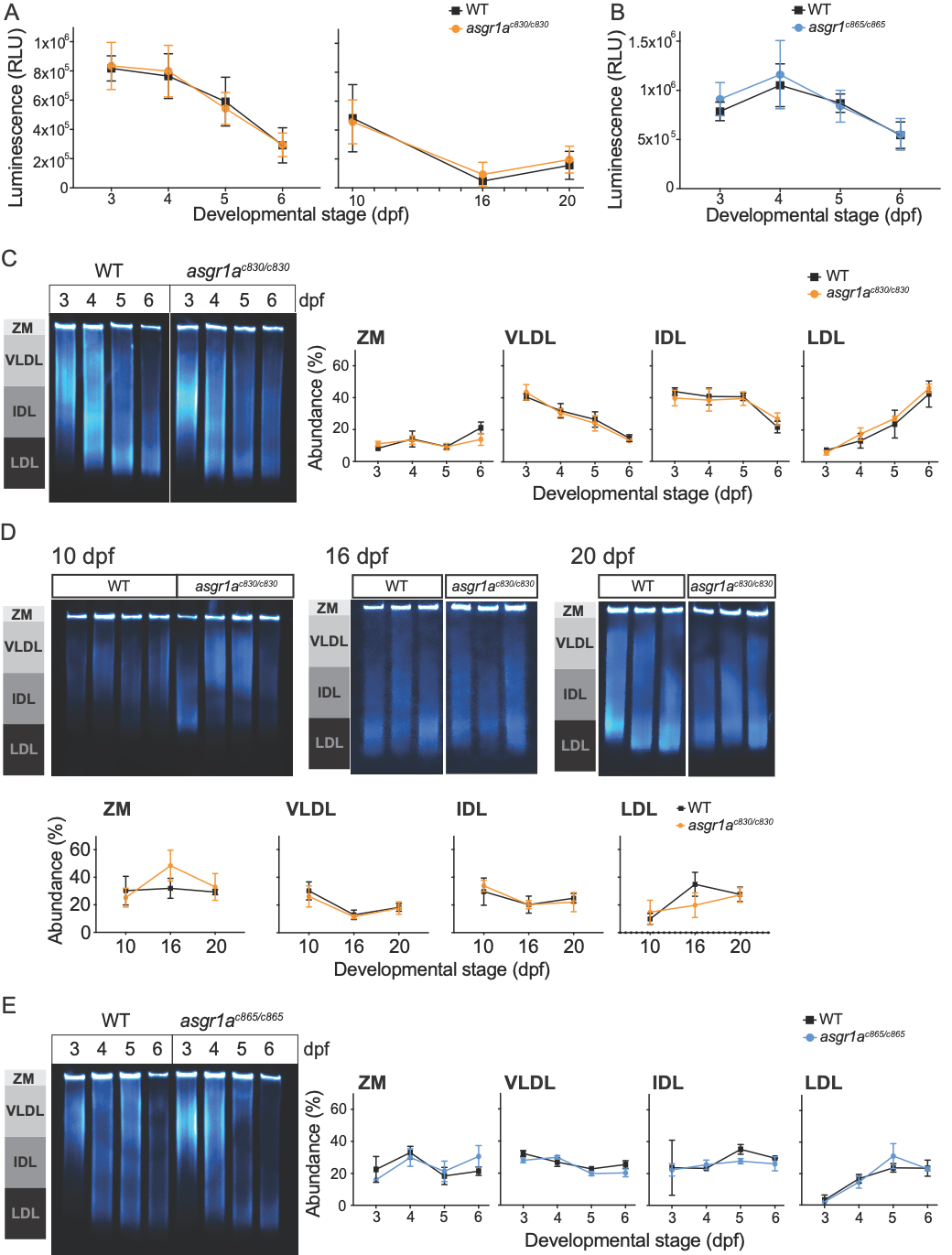
**S Figure 4: Examining B-lp quantity and particle sizes in *asgr1a* mutants.**

Endogenous fusion of NanoLuc to ApoB in the LipoGlo reporter line allows examination of B-lp levels of single larva by luminescence readout (Relative Luminescence Units: RLU). (A) LipoGlo-Counting assay of WT and *asgr1a^c830/c830^* siblings throughout larval and juvenile development. n=20-76 fish/genotype/time point pooled from 3 independent experiments. (B) LipoGlo-Counting assay of WT and *asgr1a^c865/c865^* siblings throughout larval development. n=10-107 fish/genotype/time point from 1-2 independent experiments. (C, D) Representative image and quantification of LipoGlo-Electrophoresis assay on larval (C) and juvenile (D) stages of WT and *asgr1a^c830/c830^* siblings. n=3 fish/genotype/time point representative chosen from dataset shown in (A). (E) Representative image and quantification of LipoGlo-Electrophoresis assay on larval stages of WT and *asgr1a^c865/c865^* siblings. n=3-6 fish/genotype/time point representative chosen from dataset shown in (B). No significant differences could be detected in any of LipoGlo experiments using two-way ANOVA followed by Tukey’s multiple comparison test.

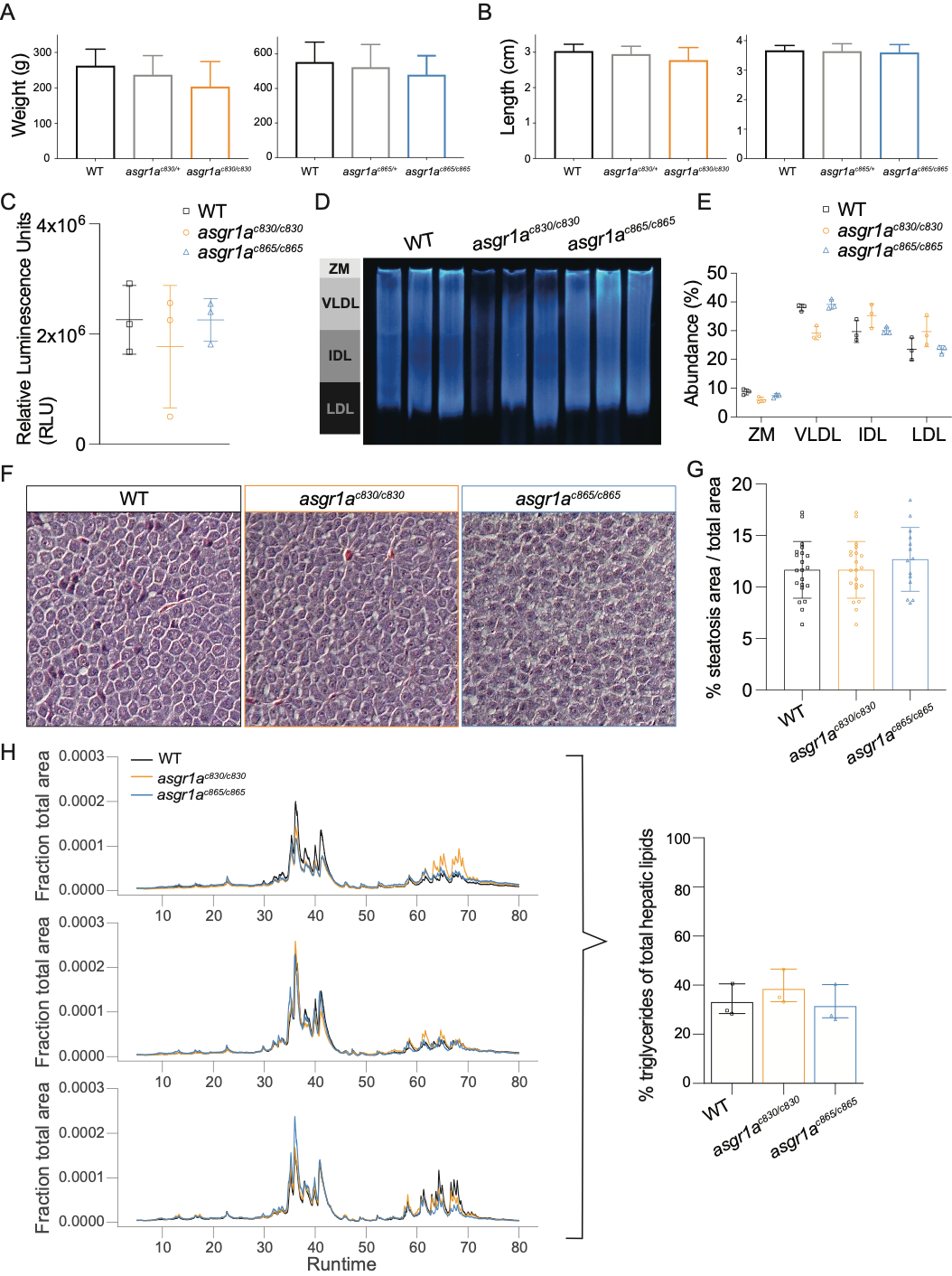

**S Figure 5: *Asgr1a* mutant adults are the same size and weight as their WT siblings and females show no phenotype after feeding a WD for 6 days.**

(A, B) Raising zebrafish from an in-cross of *asgr1a^c830/+^* and *asgr1a^c865/+^* in one tank, respectively. Hetero- and homozygous mutants are the same (A) weight and (B) length at 6 months of age. (C-E) After 6-days of WD, LipoGlo assays on blood obtained from 3 females of each mutation and controls did not show any differences in (C) B-lp quantity or particle sizes (E), representative image for LipoGlo Electrophoresis (D). (F) Representative images of liver sections from the same females as in C-E, stained with H & E. (G) Quantification of H & E images, confirming no change in steatosis levels between mutants and controls. (H) HPLC traces of three females from each genotype, with quantification not showing any changes between genotypes. n = 3 from one experiment. Analyzed using two-way ANOVA followed by Tukey’s multiple comparison test.

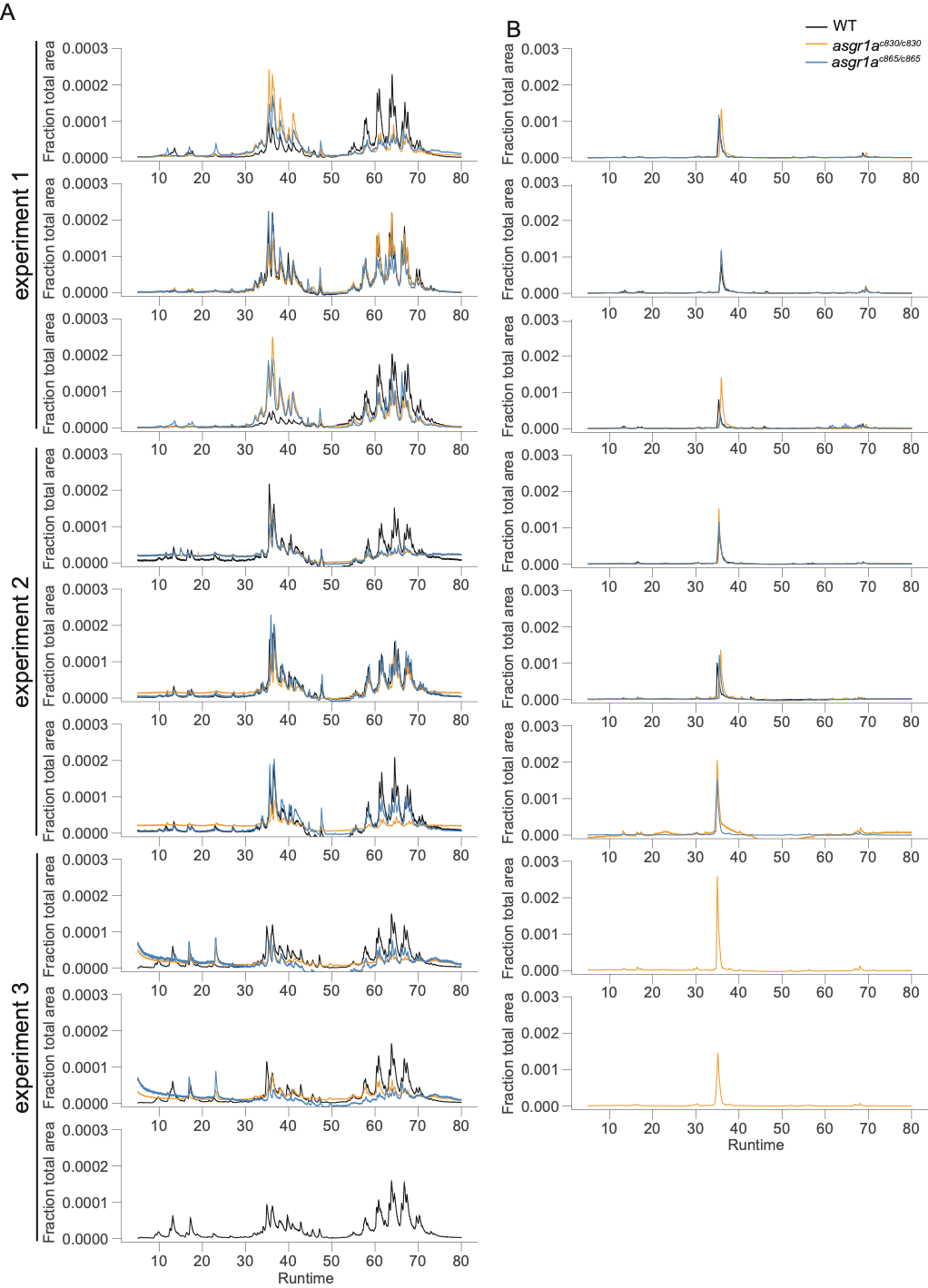

**S Figure 6: HPLC traces of all experiments performed on male zebrafish fed a WD for 6 days.**

A) HPLC traces of livers that were dissected 3 hours after the final meal on the 6th day. B) HPLC traces of male zebrafish fecal matter collected from the fish in A). Each trace represents a biological replicate, normalized to the total area of each chromatograph from 2 - 80 min. Genotypes from the same biological replicates are overlaid. Experimental replicates are grouped as shown. Insufficient mutant fish were present in the third experiment to generate three biological liver replicates, three replicates are only present for wild-type fish. Fecal samples were collected for each fish, but samples that did not pass quality control due to insufficient collected mass (peak height of cholesterol less than 10 pA) were removed from the analysis.
